# Robust inference in summary data Mendelian randomisation via the zero modal pleiotropy assumption

**DOI:** 10.1101/126102

**Authors:** Fernando Pires Hartwig, George Davey Smith, Jack Bowden

## Abstract

**Background:** Mendelian randomisation (MR) is being increasingly used to strengthen causal inference in observational studies. Availability of summary data of genetic associations for a variety of phenotypes from large genome-wide association studies (GWAS) allows straightforward application of MR using summary data methods, typically in a two-sample design. In addition to the conventional inverse variance weighting (IVW) method, recently developed summary data MR methods, such as the MR-Egger and weighted median approaches, allow a relaxation of the instrumental variable assumptions.

**Methods:** Here, a new method –the mode-based estimate (MBE) – is proposed to obtain a single causal effect estimate from multiple genetic instruments. The MBE is consistent when the largest number of similar (identical in infinite samples) individual-instrument causal effect estimates comes from valid instruments, even if the majority of instruments are invalid. We evaluate the performance of the method in simulations designed to mimic the two-sample summary data setting, and demonstrate its use by investigating the causal effect of plasma lipid fractions and urate levels on coronary heart disease risk.

**Results:** The MBE presented less bias and type-I error rates than other methods under the null in many situations. Its power to detect a causal effect was smaller compared to the IVW and weighted median methods, but was larger than that of MR-Egger regression, with sample size requirements typically smaller than those available from GWAS consortia.

**Conclusions:** The MBE relaxes the instrumental variable assumptions, and should be used in combination with other approaches in a sensitivity analysis.

**Key Messages:**

- Summary data Mendelian randomisation, typically in a two-sample setting, is being increasingly used due to the availability of summary association results from large genome-wide association studies.
- Mendelian randomisation analyses using multiple genetic instruments are prone to bias due to horizontal pleiotropy, especially when genetic instruments are selected based solely on statistical criteria.
- A causal effect estimate robust to horizontal pleiotropy can be obtained using the mode-based estimate (MBE).
- The MBE requires that the most common causal effect estimate is a consistent estimate of the true causal effect, even if the majority of instruments are invalid (i.e., the ZEro Modal Pleiotropy Assumption, or ZEMPA).
- Plotting the smoothed empirical density function is useful to explore the distribution of causal effect estimates, and to understand how the MBE is determined.

## Introduction

Using germline genetic variants as instrumental variables of modifiable exposure phenotypes can strengthen causal inference in observational studies by applying the principles of Mendelian randomisation (MR).^1,2^ This method has already been used to address causality in several exposure-outcome combinations and has become a common feature in the recent epidemiological literature. ^3^ Causal inference using MR relies on the instrumental variable assumptions, which require that the genetic variant is: i) associated with the exposure, ii) independent on confounders of the exposure-outcome associations, and iii) independent of the outcome after conditioning on the exposure and all exposure-outcome confounders.

Recent MR methods allow performing MR with multiple genetic instruments, typically single nucleotide polymorphisms (SNPs), using summary data estimates from genome-wide association studies (GWAS).^5^ Given the increasing number of publicly available summary statistics from large GWAS consortia, summary data MR methods enable many causal hypotheses to be rapidly interrogated without the administrative burden and cooperation required to perform equivalent individual-level data analyses.^6,7^

However, using many instruments in an MR analysis increases the probability of including at least one invalid instrument, which could easily bias the estimate. For example, the inverse variance weighting (IVW) method requires that either all variants are valid instruments or that there is balanced horizontal pleiotropy (i.e., horizontal pleiotropic effects of individual instruments sum to zero) and that such pleiotropic effects are independent of instrument strength across all variants (i.e., the Instrument Strength Independent of Direct Effects - InSIDE - assumption). ^5,8^ More recently, other summary data MR methods that allow relaxing (but not eliminating) the instrumental variable assumptions regarding horizontal pleiotropy have been proposed.^9,10^

In this paper, we describe a new summary data MR method - the mode-based estimate (MBE). We clarify when this will be a consistent estimate of the causal effect, compare it to established summary data MR methods using simulations, and illustrate its application using a real data example.

## Methods

In order to motivate the summary data methods discussed in this paper, we assume the following simple data generating model linking genetic variant *G*_*j*_ *(j = 1, …, L)*, a continuous exposure *X*_*i*_ and outcome *Y*_*i*_ for subject *i*:

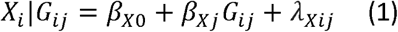

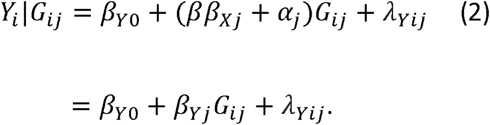

Here, *β*_*X j*_ and *β*_*Y j*_ = (*ββ*_*X j*_ + *α*_*j*_) represent *G*_*j*_’s true association with the exposure and outcome, respectively. *ββ*_*X j*_ is the effect of *G*_*j*_ on *Y* through *X*, where *β* is the causal effect of *X* on *Y* we wish to estimate. The term α_*j*_ represents the association between *G*_*j*_ and *Y* not through the exposure of interest, due to horizontal pleiotropy. The error terms *λ*_*Xij*_ and *λ*_*Yij*_ will generally be correlated when the collected on the same individuals. However, we will mainly focus on the two-sample setting where the error terms are independent, because independent samples are used to fit models (1) and (2). For simplicity, we will also assume that all L genetic variants are mutually independent of one another.

Let 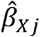 and 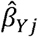 represent the SNP-exposure and SNP-outcome association estimate for variant j, respectively, and let 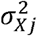 and 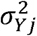 represent the variance of 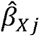 and 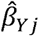, respectively. The ratio estimate^11,12^ for the causal effect *β* using variant *j* alone is equal to

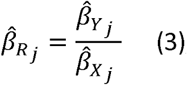

the standard error of which (*σ*_*Rj*_) can be obtained using the delta method^13^ as follows:

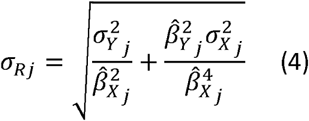

The standard error in (4) can be simplified to 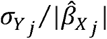 when the variance of the SNP-exposure association 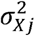 is small enough to be considered “ignorable”, or equivalently that 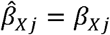. This is equivalent to the NO Measurement Error (NOME) assumption.

The ratio estimate 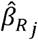 is a crude measure of causal effect, but has a major advantage over more sophisticated methods in that it can be calculated using summary data estimates for 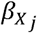 and 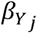 alone. These estimates can then be used furnish a summary data MR analysis using the framework of a meta-analysis.

Under models (1) and (2), variant *j* is a valid instrument when *α*_*j*_=0 and invalid when α_j_≠0. When α_j_≠0, 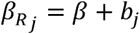 where *b*_*j*_ = α_j_/β_*x j*_a bias term). In the Supplementary Methods, we briefly review three such summary data methods – IVW^5^, MR-Egger regression^9^ and weighted median^10^ - and discuss the conditions under which each method returns a consistent causal effect estimate (i.e., estimate converges in probability to the true value as the sample size increases).

### The MBE

In this paper we propose a new causal effect estimator - the MBE - that offers robustness to horizontal pleiotropy in a different manner to that of the IVW, MR-Egger or weighted median methods. Its ability to consistently estimate the true causal effect relies on the following fundamental assumption termed the ZEro Modal Pleiotropy Assumption (ZEMPA):

ZEMPA: *Across all instruments, the most frequent value (i.e., the mode) of b*_*j*_ *is 0.*

In order to formalize this, let *k ∈* {1,2,……, *L*} represent the number of unique values of *b*_*j*_ among the *L* variants. If all *b*_*j*_ terms are identical then *k*=1, but if all are unique then *k= L.* Now, let *n*_1_,*n*_2_,…, *n*_*k*_ represent the number of instruments that have the same non-zero value of *b*_*j*_, where *n*_1_ represents those with the smallest non-zero identical value of *b*_*j*_ and *n*_*k*_ represents those with the largest non-zero identical value. Finally, let *n*_0_ represent the number of valid instruments whose *b*_*j*_ terms are identically zero. We then have that *n*_0_ + *n*_1_ + … + *n*_*k*_ = *L*. ZEMPA implies that *n*_0_ is larger than any other *n*_*l*_ for *l* in 1, 2,…, *k* (i.e., *n*_0_ > Max(*n*_1_,…, *n*_*k*_)). For a weighted MBE, that is an MBE derived by allowing the weight given to each ratio estimate to vary, ZEMPA implies that the weights associated with the valid instruments are the largest among all *k* subsets of instruments (ie., *w*_*0*_ > Max(*w*_*1*_,…, *w*_*k*_), where *w*_*l*_ is the weight contributed by the *l*th subset of instruments using our previous subset definition based on *b*_*j*_.

The breakdown level (i.e., the maximum proportion of information that can come from invalid instruments before the method is inconsistent) of the simple MBE ranges from 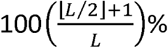 to 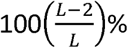. The lower limit corresponds to the situation where there are some valid instruments, but all invalid instruments estimate the same (biased) causal effect parameter (ie, *k*= 2) implying that ZEMPA is satisfied (ie, *n*_0_ > Max(*n*_1_,…, *n*_*k*_)) if up to, but not including, half of the instruments are invalid. The upper limit corresponds to the situation where all invalid instruments estimate different causal effect parameters (ie, *n*_1_ = *n*_2_ = … = *n*_*k*_ = 1), implying that ZEMPA would be satisfied if just two variants were valid (*n*_0_ = 2) and the remainder (*L – 2*) were invalid. Given that Max(*n*_1_,…, *n*_*k*_) is often unknown and is likely to vary depending on the set of genetic instruments and the outcome variable, the true breakdown level of the MBE in any given applied investigation is difficult to determine.

For example, in Figure 1A, 6 out of 8 instruments are invalid (so *n*_0_ = 2), but all non-zero *b*_*j*_s are unique, implying that *k= L* – 1 = 7 and *n*_1_ = *n*_2_ = … = *n*_7_ = 1. In this situation, ZEMPA is satisfied and the simple MBE is a consistent estimate of the causal effect *β*. However, when the largest number of identical estimates comes from invalid instruments (i.e., *n*_0_ < *n*_*l*_ for some *l*, ZEMPA violated), then the simple MBE will be inconsistent for *β* (i.e., asymptotically biased). This is illustrated in Figure 1B, which shows causal effect estimates from 6 invalid and 2 valid variants (*n*_0_ = 2). Since three variants have precisely the same horizontal pleiotropic effect in this example (*n*_2_ = 3), ZEMPA is violated.

**Figure 1.**
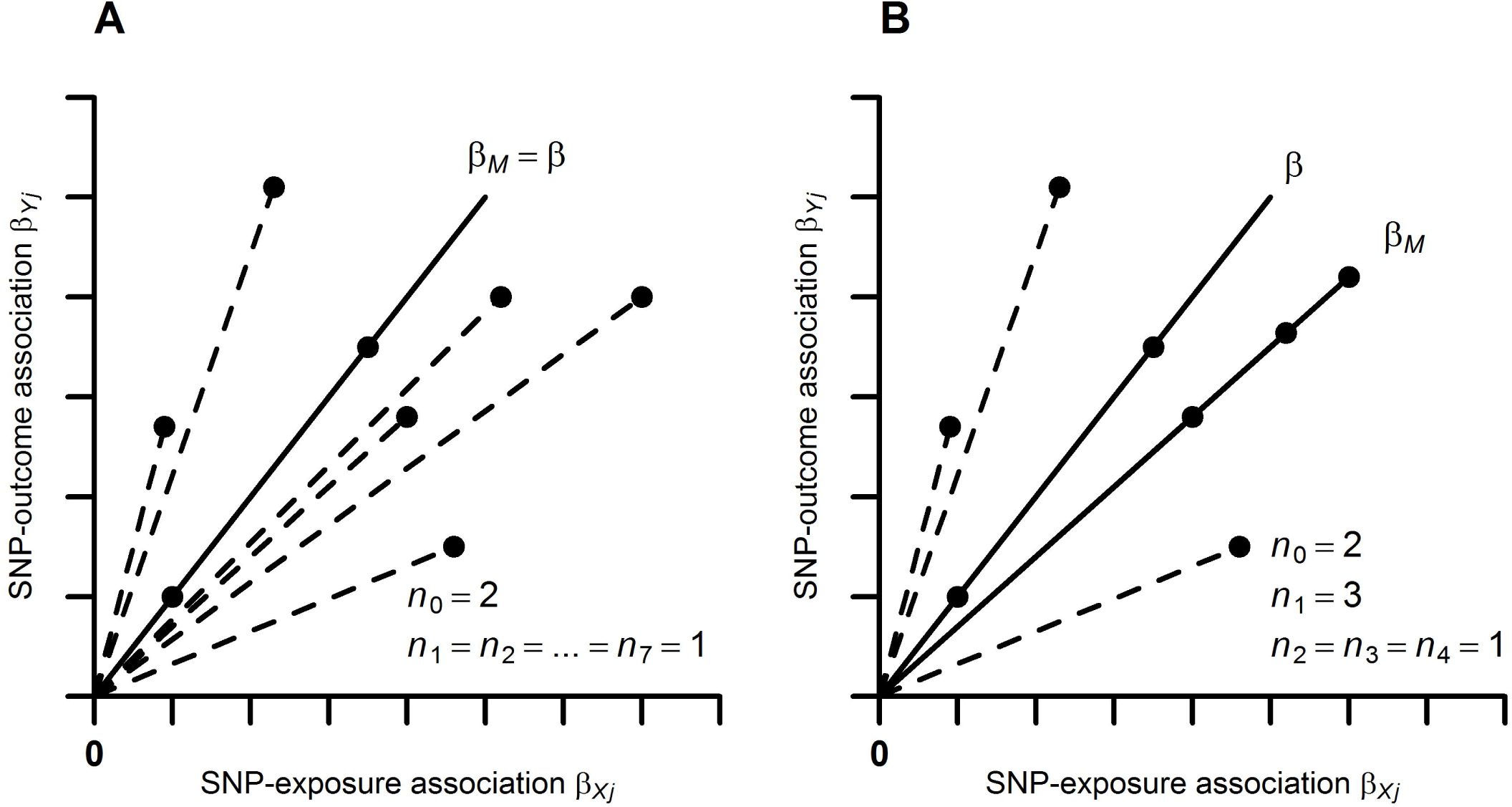
Illustration of the ZEro Modal Pleiotropy Assumption (ZEMPA) in the simple mode-based estimate (MBE). *β*_*M*_ is the simple MBE causal effect and *β* is the true causal effect. *n*_*l*_ denotes the number of variants with a given horizontal pleiotropic effect (*n*_0_ denotes the number of valid instruments). Panel A: ZEMPA is satisfied. Panel B: ZEMPA is violated. SNP: Single nucleotide polymorphism.

The breakdown level of the weighted MBE can be similarly defined as ranging from 50% (exclusive) to 100% (exclusive). In other words, the weighted MBE is biased if *w*_0_ < *w*_*l*_ for some *l*. Of note, the limits are open intervals because the weights are real numbers, unlike number of instruments (in the case of the simple MBE), which is a natural number. However, as *L* increases, then the lower and upper limits of the breakdown level of the simple MBE also tend to 50% and 100%, respectively.

### Implementing the MBE

To calculate the MBE, we propose using the mode of the smoothed empirical density function of all 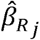 as the causal effect estimate. This strategy is straightforward to implement, easily deals with sampling variation in asymptotically identical 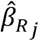s and allows giving different weights to different instruments. We refer to the mode of the unweighted and inverse-variance weighted empirical density function as the simple and weighted MBE’s, respectively. The standardised weights for the weighted MBE can be computed as follows:

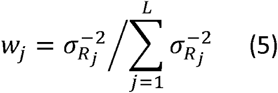

For the simple MBE, *w*_1_ = *w*_2_ = = *w*_L_ = 1/*L*.

Consider the normal kernel density function of the 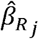:

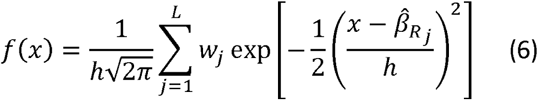

where *h* is the smoothing bandwidth parameter.^16^ The causal effect estimate obtained using the MBE method 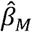 is the value of *x* that maximizes *f(x)*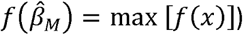. The *h* parameter regulates a bias-variance trade-off of the MBE, with increasing *h* leading to higher precision, but also to higher bias. Here, *h* = *φs*, with *φ* being a tuning parameter that allows increasing or decreasing the bandwidth, and *s* being the default bandwidth value chosen according to some criterion. We used the modified Silverman’s bandwidth rule proposed by Bickel^17^:

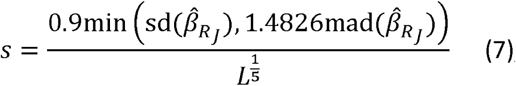

where sd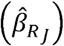 and mad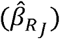 are the standard deviation and median absolute deviation from the median of the 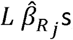, respectively.

An intuitive explanation of the MBE based on an analogy with histograms is provided in the Supplementary Methods.

### Simulation model

The simulations were performed using the following model to generate individual *i*’s exposure *x*_*i*_, outcome *Y*_*i*_ and confounder *U*_*i*_ based on their underlying genetic data vector (*G*_*i1*_,…,*G*_*iL*_):

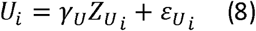

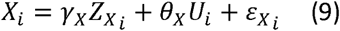

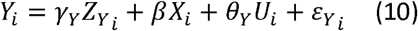

where:

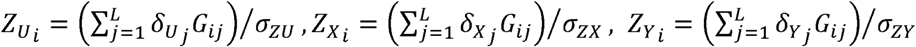

*Z*_*U*_, *Z*_X_ and *Z*_*Y*_ represent the additive allele scores of *L* independent SNPs on *U*, *X* and *Y*, modulated by the parameters 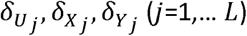. *β* denotes the true causal effect of *X* on *Y* that we wish to estimate. The underlying genetic variables (*G*_*ij*_) were generated independently by sampling from a Binomial (2, *p*) distribution with *p* itself drawn from a Uniform(0.1,0.9) distribution, to mimic bi-allelic SNPs in Hardy-Weinberg equilibrium. The resulting allele scores were then divided by their sample standard deviations (*σ*_*ZU*_, *σ*_*ZX*_, *σ*_*ZY*_), to set variances to one. The direct effects of *U* on *X* and *Y* are denoted by *θ*_*X*_ and *θ*_*Y*_, respectively. *θ*_*X*_ and *θ*_*Y*_ are set to positive values in all simulations, so as to always induce positive confounding. Error terms 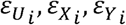, were independently generated from a normal distribution, with mean=0 and variances, 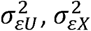 and 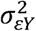 respectively, whose values were chosen to set the variances of *U*, *X* and *Y* to one.

Constraining the variances in this way enables easy interpretation of the parameters in models (8)-(10). For example, *β* =0.1 implies that one standard deviation increment in *X* causes a 0.1 standard deviation increment in *Y*, and that the causal effect of *X* on *Y* explains 1% of *Y* variance.

A summary data interpretation of our simulation model is provided in the Supplementary Methods.

### Simulation scenarios

Although the consistency property of an estimator provides a formal justification of the approach, it is equally important to understand how well it works in practice for realistically sized data sets in comparison to other methods. Therefore, we evaluated our proposed estimator in four different simulation scenarios. In all simulations the number of variants *L*=30, 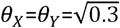, 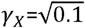 and 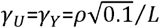, where *ρ* = 0, 3, 6, …, 30 is the number of invalid instruments.

Simulations 1 and 2 were aimed at evaluating the performance of the MBE under the causal null (*β*=0) in the two-sample setting. Datasets of 100,000 individuals were simulated and divided in half at random, and each was used to estimate either SNP-exposure or SNP-outcome associations. Simulations 3 and 4 were aimed at evaluating weak instrument bias in the two-sample and single-sample settings; sample sizes used to estimate instrument-exposure (*N*_*X*_) and instrument-outcome (*N*_*Y*_) associations were allowed to vary, as described below.

*Simulation 1:* In this scenario, 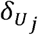 was 0 for all instruments, implying that there is no InSIDE-violating horizontal pleiotropy. InSIDE-respecting horizontal pleiotropic effects 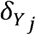 were drawn from a Uniform(0.01, 0.2) distribution for the *ρ* invalid instruments or were set to 0 for valid instruments. Moreover, given that *β*=0, power can be interpreted as the type-I error rate.

*Simulation 2:* InSIDE-violating horizontal pleiotropy was induced by setting 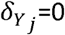 for all instruments, while 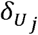 values were drawn from a Uniform(0.01, 0.2) distribution for the *ρ* invalid instruments.

*Simulation 3:* This simulation evaluated the performance of the estimators to detect a positive causal effect of *β*=0.1 in the two-sample context. 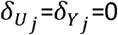 for all L instruments, and *ρ* =0, implying that there is no horizontal pleiotropy. *N*_*X*_ ∈ {25,000, 50,000, 100,000}, and *N*_*Y*_ ∈ {25,000, 50,000, 100,000}.

*Simulation 4:* This simulation evaluated the performance of the estimators under the causal null when SNP-exposure and SNP-outcome associations are estimated in partially (50%) or fully (100%) overlapping samples (the latter being equivalent to the single sample setting). It was implemented as for simulation 3, except *β*=0 and *N*_*X*_ = *N*_*Y*_ ∈ {1,000, 5,000, 10,000}. We used smaller sample sizes to purposely increase the bias due to sample overlap, thus facilitating comparisons between methods.

### Applied examples: plasma lipid fractions and urate levels and coronary heart disease risk

Do and colleagues^18^ performed a two-sample MR analysis to evaluate the causal effect of low-density lipoprotein cholesterol (LDL-C), high-density lipoprotein cholesterol (HDL-C) and triglycerides on coronary heart disease (CHD) risk, using a total of 185 genetics variants. Summary association results were obtained from the Global Lipids Genetics Consortium^19^ and the Coronary Artery Disease Genome-Wide Replication and Meta-Analysis Consortium^20^, and were downloaded from Do and colleagues’ supplementary material (standard errors were estimated based on the regression coefficients and P-values). Genetic variants were classified as instruments for each lipid fraction using a statistical criterion (P<1x10^−8^), resulting in 73 instruments for LDL-C, 85 for HDL-C and 31 for triglycerides.

White and colleagues^21^ performed a similar analysis, but with plasma urate levels rather than lipid fractions. 31 variants associated with urate levels (P<5x10^−7^) and the required summary statistics were obtained from the GWAS catalog (https://www.ebi.ac.uk/gwas/).

### Statistical analyses

In all simulation scenarios, causal effect estimates were obtained using established MR methods (multiplicative random effects IVW^8^, multiplicative random effects MR-Egger regression^8^ and weighted median, all implemented using inverse-variance weights calculated under NOME), as well as the simple and the weighted MBEs. Each version of the MBE was evaluated using weights calculated with and without making the NOME assumption (see earlier discussion), thus yielding four MBE’s. Each of these four methods was evaluated for two values of the tuning parameter Ψ ∈ {1,0.5}, totalling eight versions of the MBE method. Parametric bootstrap was used to estimate the standard errors of the MBE using the median absolute deviation from the median of the bootstrap distribution of causal effect estimates. These were used to derive symmetric confidence intervals, after multiplying by 1.4826 for asymptotically normal consistency.

In each scenario, coverage, power and average causal effect estimates, standard errors, 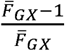 and 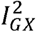 statistics (which quantify the magnitude of violation of the NOME assumption in IVW and MR-Egger regression estimates, respectively^8,14^) were obtained across 10,000 simulated datasets. Power was defined as the proportion of times that 95% confidence intervals excluded zero, and coverage as the proportion of times that 95% confidence intervals included the true causal effect.

MR methods were also applied to estimate the causal effect of plasma lipid fractions and urate levels on CHD risk. The magnitude of regression dilution bias in IVW and MR-Egger regression was assessed by the 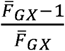 and 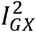 statistics, respectively. Cochran’s Q test was used to test for the presence of and horizontal pleiotropy (under the assumption that this is the only source of heterogeneity between 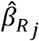 other than chance).^15^ All simulations and analyses were performed using R 3.3.1 (www.r-project.org). R code for implementing the MBE is provided in the Supplementary Methods.

## Results

### Performance under the causal null in the two-sample context

The results of simulation 1 - where directional horizontal pleiotropy (if any) occurs only under the InSIDE assumption - are shown in Table 1. When all instruments are valid, all methods were unbiased with type-I error rates ≤5%. As expected, MR-Egger regression (which is consistent if InSIDE holds) was the least biased method in this scenario, especially when many instruments were invalid. The four MBE’s in Table 1 were less biased and less precise than the IVW and the weighted median methods. The simple MBE was more biased than the weighted MBE (noticeable especially when the proportion of invalid instruments was high). Using weights derived under the NOME assumption increased bias and false rejection rates. Setting *φ*=0.5 (i.e., setting the bandwidth to half of the default value) reduced both bias and precision (Supplementary Table 1).

When InSIDE is violated (Table 2), again the MBE’s were less biased than IVW and weighted median methods. In this case, however, they were also less biased than MR-Egger regression estimates, which is known to be highly sensitive to InSIDE violation^8^. The exception was for large proportions (i.e., ≥80%) of invalid instruments, where MR-Egger estimates were the least biased. This is because the degree of InSIDE violation, as quantified by an inverse variance weighted Pearson correlation between instrument strength and horizontal pleiotropic effects^8^, is smaller in those situations (Supplementary Table 3). Moreover, in this scenario, the simple MBE was generally less biased than their weighted counterparts, and setting *φ*=0.5 had a smaller effect when compared to simulation 1 (and indeed only clear for the simple MBE - Supplementary Table 3). The NOME assumption again increased bias and false rejection rates.

**Table 1.** Mean estimates from simulation 1: directional horizontal pleiotropy under the lnSlDE assumption and zero causal effect (10,000 simulations per scenario).

| Estimator | Statistic | Proportion (%) of invalid instruments (Mean $\frac{\bar{F}_{GX}-1}{\bar{F}_{GX}}$ [%]; mean $I_{GX}^2$ [%]) | | | | | | | | | | |
| --- | --- | --- | --- | --- | --- | --- | --- | --- | --- | --- | --- | --- |
|  |  | 0<br>(99.7;<br>97.4) | 10<br>(99.7;<br>97.4) | 20<br>(99.7;<br>97.4) | 30<br>(99.7;<br>97.4) | 40<br>(99.7;<br>97.4) | 50<br>(99.7;<br>97.4) | 60<br>(99.7;<br>97.4) | 70<br>(99.7;<br>97.4) | 80<br>(99.7;<br>97.4) | 90<br>(99.7;<br>97.4) | 100<br>(99.7;<br>97.4) |
| IVW | Beta | 0.000 | 0.081 | 0.159 | 0.238 | 0.315 | 0.394 | 0.473 | 0.550 | 0.629 | 0.707 | 0.784 |
|  | SE | 0.015 | 0.058 | 0.078 | 0.092 | 0.102 | 0.109 | 0.114 | 0.117 | 0.118 | 0.117 | 0.115 |
|  | Coverage (%) | 96.9 | 86.7 | 51.1 | 18.9 | 4.8 | 0.6 | 0.1 | 0.0 | 0.0 | 0.0 | 0.0 |
|  | Power (%)<sup>a</sup> | 3.2 | 13.3 | 48.9 | 81.1 | 95.2 | 99.4 | 99.9 | 100.0 | 100.0 | 100.0 | 100.0 |
| MR-Egger | Beta | 0.001 | 0.003 | 0.006 | 0.008 | 0.004 | 0.010 | 0.014 | 0.011 | 0.020 | 0.018 | 0.018 |
|  | SE | 0.032 | 0.127 | 0.170 | 0.197 | 0.215 | 0.226 | 0.231 | 0.230 | 0.224 | 0.212 | 0.191 |
|  | Coverage (%) | 96.6 | 95.5 | 94.7 | 94.3 | 94.2 | 93.9 | 94.0 | 93.9 | 94.0 | 93.6 | 93.5 |
|  | Power (%)<sup>a</sup> | 3.4 | 4.5 | 5.3 | 5.7 | 5.8 | 6.1 | 6.0 | 6.1 | 6.1 | 6.4 | 6.5 |
| Weighted<br>Median | Beta | 0.000 | 0.008 | 0.020 | 0.037 | 0.076 | 0.168 | 0.305 | 0.433 | 0.541 | 0.624 | 0.692 |
|  | SE | 0.020 | 0.021 | 0.023 | 0.026 | 0.031 | 0.038 | 0.043 | 0.044 | 0.044 | 0.043 | 0.043 |
|  | Coverage (%) | 97.6 | 95.6 | 88.1 | 73.3 | 48.7 | 20.4 | 4.7 | 0.5 | 0.0 | 0.0 | 0.0 |
|  | Power (%)<sup>a</sup> | 2.5 | 4.4 | 11.9 | 26.7 | 51.3 | 79.6 | 95.3 | 99.5 | 100.0 | 100.0 | 100.0 |
| Simple<br>MBE<sup>b</sup> | Beta | 0.000 | 0.000 | 0.002 | 0.003 | 0.013 | 0.040 | 0.129 | 0.294 | 0.501 | 0.650 | 0.742 |
|  | SE | 0.046 | 0.054 | 0.056 | 0.069 | 0.084 | 0.118 | 0.126 | 0.148 | 0.183 | 0.175 | 0.183 |
|  | Coverage (%) | 99.2 | 98.8 | 98.5 | 97.9 | 96.8 | 87.0 | 37.4 | 9.9 | 5.6 | 4.4 | 4.1 |
|  | Power (%)<sup>a</sup> | 0.8 | 1.2 | 1.5 | 2.1 | 3.2 | 13.0 | 62.6 | 90.1 | 94.4 | 95.6 | 95.9 |
| Weighted<br>MBE<sup>b</sup> | Beta | 0.000 | 0.001 | 0.001 | 0.003 | 0.014 | 0.044 | 0.125 | 0.222 | 0.332 | 0.430 | 0.513 |
|  | SE | 0.040 | 0.048 | 0.050 | 0.063 | 0.076 | 0.107 | 0.103 | 0.107 | 0.135 | 0.132 | 0.144 |
|  | Coverage (%) | 98.5 | 98.0 | 97.6 | 96.6 | 93.8 | 71.1 | 19.5 | 8.2 | 6.8 | 5.6 | 5.1 |
|  | Power (%)<sup>a</sup> | 1.5 | 2.0 | 2.4 | 3.4 | 6.2 | 28.9 | 80.5 | 91.8 | 93.3 | 94.4 | 94.9 |
| Simple<br>MBE<br>(Under<br>NOME)<sup>b</sup> | Beta | 0.000 | 0.000 | 0.002 | 0.003 | 0.013 | 0.040 | 0.129 | 0.294 | 0.501 | 0.650 | 0.742 |
|  | SE | 0.032 | 0.031 | 0.031 | 0.032 | 0.035 | 0.043 | 0.054 | 0.071 | 0.076 | 0.070 | 0.066 |
|  | Coverage (%) | 99.1 | 98.7 | 98.1 | 97.4 | 95.8 | 84.7 | 29.8 | 4.1 | 1.4 | 0.6 | 0.6 |
|  | Power (%)<sup>a</sup> | 0.9 | 1.3 | 1.9 | 2.6 | 4.3 | 15.4 | 70.2 | 96.0 | 98.6 | 99.4 | 99.4 |
| Weighted<br>MBE<br>(Under<br>NOME)<sup>b</sup> | Beta | 0.000 | 0.001 | 0.002 | 0.004 | 0.016 | 0.063 | 0.193 | 0.343 | 0.481 | 0.577 | 0.644 |
|  | SE | 0.026 | 0.026 | 0.026 | 0.026 | 0.029 | 0.039 | 0.045 | 0.049 | 0.049 | 0.045 | 0.045 |
|  | Coverage (%) | 98.3 | 97.6 | 97.2 | 95.8 | 92.3 | 65.8 | 12.0 | 2.3 | 1.2 | 0.7 | 0.8 |
|  | Power (%)<sup>a</sup> | 1.7 | 2.4 | 2.9 | 4.2 | 7.7 | 34.2 | 88.0 | 97.8 | 98.8 | 99.3 | 99.2 |

**Table 2.**
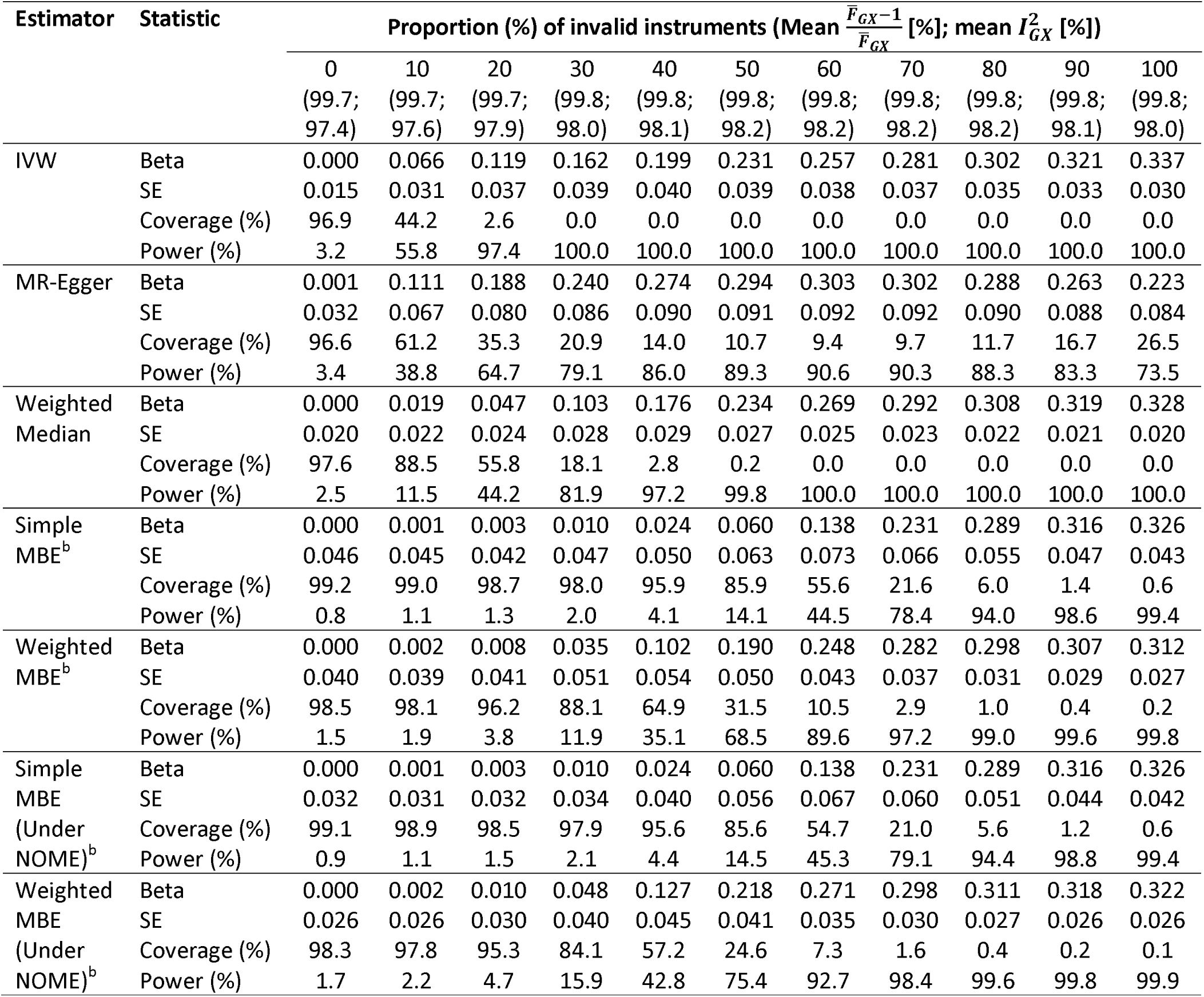
Mean estimates from simulation 2: directional horizontal pleiotropy mediated by a single confounder of the exposure-outcome association (so violating the lnSlDE assumption) and zero causal effect (10,000 simulations per scenario).

| Estimator | Statistic | Proportion (%) of invalid instruments (Mean $\frac{\bar{F}_{GX}-1}{\bar{F}_{GX}}$ [%]; mean $I_{GX}^2$ [%]) | | | | | | | | | | |
| --- | --- | --- | --- | --- | --- | --- | --- | --- | --- | --- | --- | --- |
|  |  | 0<br>(99.7;<br>97.4) | 10<br>(99.7;<br>97.6) | 20<br>(99.7;<br>97.9) | 30<br>(99.8;<br>98.0) | 40<br>(99.8;<br>98.1) | 50<br>(99.8;<br>98.2) | 60<br>(99.8;<br>98.2) | 70<br>(99.8;<br>98.2) | 80<br>(99.8;<br>98.2) | 90<br>(99.8;<br>98.1) | 100<br>(99.8;<br>98.0) |
| IVW | Beta | 0.000 | 0.066 | 0.119 | 0.162 | 0.199 | 0.231 | 0.257 | 0.281 | 0.302 | 0.321 | 0.337 |
|  | SE | 0.015 | 0.031 | 0.037 | 0.039 | 0.040 | 0.039 | 0.038 | 0.037 | 0.035 | 0.033 | 0.030 |
|  | Coverage (%) | 96.9 | 44.2 | 2.6 | 0.0 | 0.0 | 0.0 | 0.0 | 0.0 | 0.0 | 0.0 | 0.0 |
|  | Power (%) | 3.2 | 55.8 | 97.4 | 100.0 | 100.0 | 100.0 | 100.0 | 100.0 | 100.0 | 100.0 | 100.0 |
| MR-Egger | Beta | 0.001 | 0.111 | 0.188 | 0.240 | 0.274 | 0.294 | 0.303 | 0.302 | 0.288 | 0.263 | 0.223 |
|  | SE | 0.032 | 0.067 | 0.080 | 0.086 | 0.090 | 0.091 | 0.092 | 0.092 | 0.090 | 0.088 | 0.084 |
|  | Coverage (%) | 96.6 | 61.2 | 35.3 | 20.9 | 14.0 | 10.7 | 9.4 | 9.7 | 11.7 | 16.7 | 26.5 |
|  | Power (%) | 3.4 | 38.8 | 64.7 | 79.1 | 86.0 | 89.3 | 90.6 | 90.3 | 88.3 | 83.3 | 73.5 |
| Weighted<br>Median | Beta | 0.000 | 0.019 | 0.047 | 0.103 | 0.176 | 0.234 | 0.269 | 0.292 | 0.308 | 0.319 | 0.328 |
|  | SE | 0.020 | 0.022 | 0.024 | 0.028 | 0.029 | 0.027 | 0.025 | 0.023 | 0.022 | 0.021 | 0.020 |
|  | Coverage (%) | 97.6 | 88.5 | 55.8 | 18.1 | 2.8 | 0.2 | 0.0 | 0.0 | 0.0 | 0.0 | 0.0 |
|  | Power (%) | 2.5 | 11.5 | 44.2 | 81.9 | 97.2 | 99.8 | 100.0 | 100.0 | 100.0 | 100.0 | 100.0 |
| Simple<br>MBE<sup>b</sup> | Beta | 0.000 | 0.001 | 0.003 | 0.010 | 0.024 | 0.060 | 0.138 | 0.231 | 0.289 | 0.316 | 0.326 |
|  | SE | 0.046 | 0.045 | 0.042 | 0.047 | 0.050 | 0.063 | 0.073 | 0.066 | 0.055 | 0.047 | 0.043 |
|  | Coverage (%) | 99.2 | 99.0 | 98.7 | 98.0 | 95.9 | 85.9 | 55.6 | 21.6 | 6.0 | 1.4 | 0.6 |
|  | Power (%) | 0.8 | 1.1 | 1.3 | 2.0 | 4.1 | 14.1 | 44.5 | 78.4 | 94.0 | 98.6 | 99.4 |
| Weighted<br>MBE<sup>b</sup> | Beta | 0.000 | 0.002 | 0.008 | 0.035 | 0.102 | 0.190 | 0.248 | 0.282 | 0.298 | 0.307 | 0.312 |
|  | SE | 0.040 | 0.039 | 0.041 | 0.051 | 0.054 | 0.050 | 0.043 | 0.037 | 0.031 | 0.029 | 0.027 |
|  | Coverage (%) | 98.5 | 98.1 | 96.2 | 88.1 | 64.9 | 31.5 | 10.5 | 2.9 | 1.0 | 0.4 | 0.2 |
|  | Power (%) | 1.5 | 1.9 | 3.8 | 11.9 | 35.1 | 68.5 | 89.6 | 97.2 | 99.0 | 99.6 | 99.8 |
| Simple<br>MBE<br>(Under<br>NOME)<sup>b</sup> | Beta | 0.000 | 0.001 | 0.003 | 0.010 | 0.024 | 0.060 | 0.138 | 0.231 | 0.289 | 0.316 | 0.326 |
|  | SE | 0.032 | 0.031 | 0.032 | 0.034 | 0.040 | 0.056 | 0.067 | 0.060 | 0.051 | 0.044 | 0.042 |
|  | Coverage (%) | 99.1 | 98.9 | 98.5 | 97.9 | 95.6 | 85.6 | 54.7 | 21.0 | 5.6 | 1.2 | 0.6 |
|  | Power (%) | 0.9 | 1.1 | 1.5 | 2.1 | 4.4 | 14.5 | 45.3 | 79.1 | 94.4 | 98.8 | 99.4 |
| Weighted<br>MBE<br>(Under<br>NOME)<sup>b</sup> | Beta | 0.000 | 0.002 | 0.010 | 0.048 | 0.127 | 0.218 | 0.271 | 0.298 | 0.311 | 0.318 | 0.322 |
|  | SE | 0.026 | 0.026 | 0.030 | 0.040 | 0.045 | 0.041 | 0.035 | 0.030 | 0.027 | 0.026 | 0.026 |
|  | Coverage (%) | 98.3 | 97.8 | 95.3 | 84.1 | 57.2 | 24.6 | 7.3 | 1.6 | 0.4 | 0.2 | 0.1 |
|  | Power (%) | 1.7 | 2.2 | 4.7 | 15.9 | 42.8 | 75.4 | 92.7 | 98.4 | 99.6 | 99.8 | 99.9 |

### Power to detect a causal effect in the two-sample context

Table 3 displays the results for simulation 3 (no invalid instruments). The IVW method was the most powered to detect a causal effect, followed by the weighted median method, the weighted MBE, the simple MBE and MR-Egger regression. Assuming NOME reduced the bias towards the null in the weighted MBE’s and improved power. Setting *φ*=0.5 had no consistent effect on bias, but substantially reduced power (Supplementary Table 4).

**Table 3.**
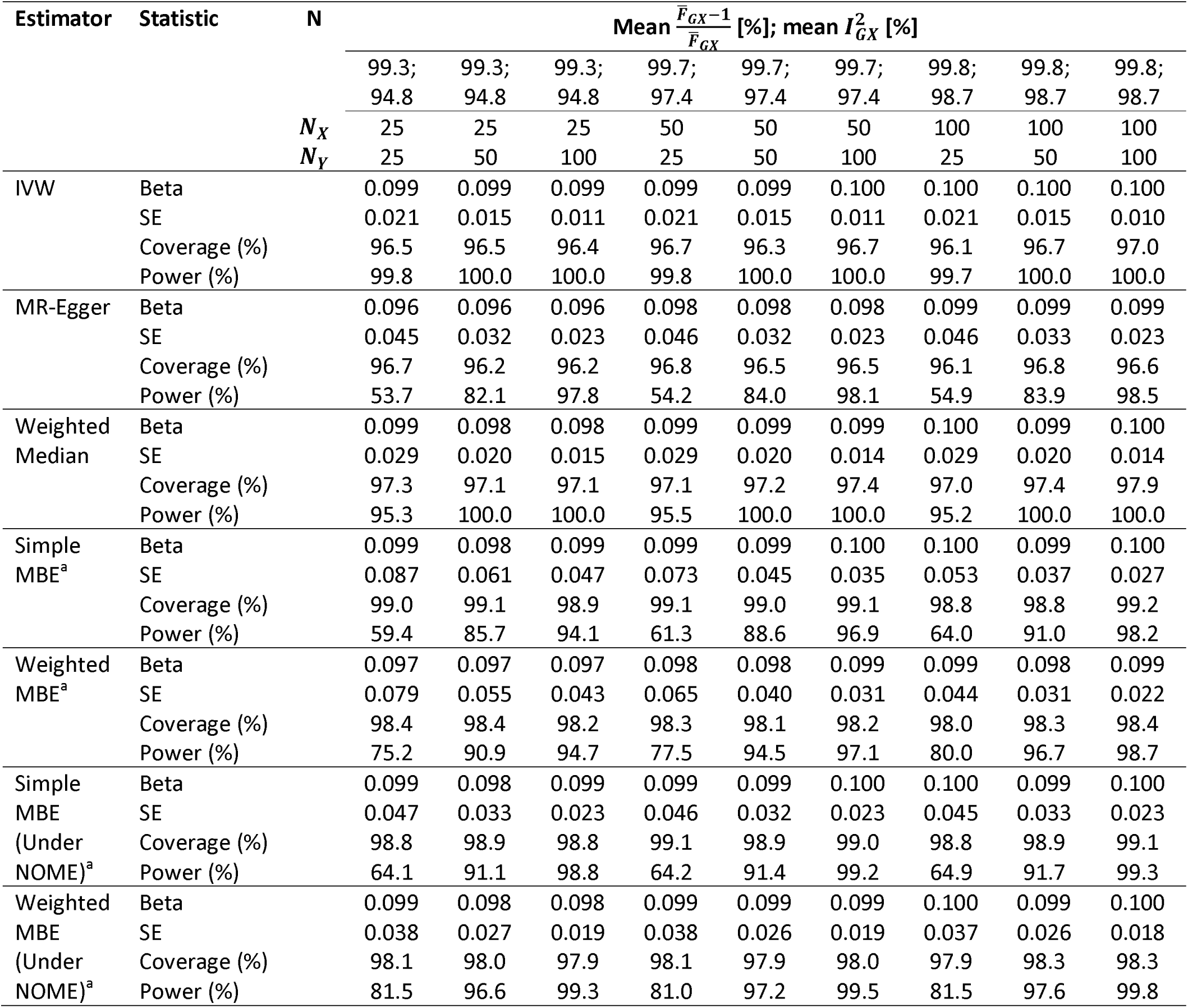
Mean estimates from simulation 3: no horizontal pleiotropy and causal effect *β* =0.1 (10,000 simulations per scenario). Sample sizes *N*_*X*_ and N_*y*_ are in thousands.

**Performance under the causal null in overlapping samples**

Supplementary Table 5 displays the performance of the methods under the causal null when the samples used to estimate instrument-exposure and instrument-outcome associations overlap. MR-Egger regression presented the largest bias, followed by the weighted MBE assuming NOME, the weighted MBE not assuming NOME, weighted median, simple MBE and IVW. Setting *φ*=0.5 slightly increased the bias (Supplementary Table 6). Importantly, the precision of the MBE was very low, suggesting that the method may be prohibitively underpowered in small samples, thus being best suited for the two-sample setting using precise summary association results. Gains in precision by making the NOME assumption was more noticeable than in the other simulations with larger sample sizes.

### Causal effect of plasma lipid fractions and urate levels on CHD risk

We used these datasets of summary association results to further explore the influence of the *φ* parameter on the MBE. First, we visually explored the distribution of ratio estimates (Figure 2). In the case of LDL-C (panel A), most of the distribution was above zero, and increasing the stringency of *φ* did not reveal substantial multimodality, although there were some pronounced density peaks at the left of the main distribution (which corresponds to the true causal effect under the ZEMPA assumption), which may result in attenuation of the causal effect estimate. However, setting *φ* =0.25 resulted in some small peaks in the main distribution which may suggest over-stringency, so we used *φ* =0.5 in the MR analysis. For HDL-C (panel B), the bulk of the distribution was centred close to zero, and setting *φ* =0.25 revealed some peaks at the left of the main distribution, suggesting that horizontal pleiotropy could lead to an apparent protective effect. Since setting *φ* =0.5 was sufficient to substantially reduce the density at the tails, this was used in the MR analysis. Regarding triglycerides (panel C), the main distribution was above zero, and the plot suggested that there may be negative horizontal pleiotropy, leading to an underestimation of the causal effect (*φ* =0.25 was used in MR analysis). Finally, in the case of urate levels (panel D), by decreasing *φ* it became increasingly evident that the distribution of was bi-modal, which could only be clearly distinguished by setting *φ* =0.25 (which was used in MR analysis) because they were similar to one another. Comparing the two distributions, the main one was the closest to zero, suggesting that horizontal pleiotropy is biasing the causal effect estimate upwards.

**Figure 2.**
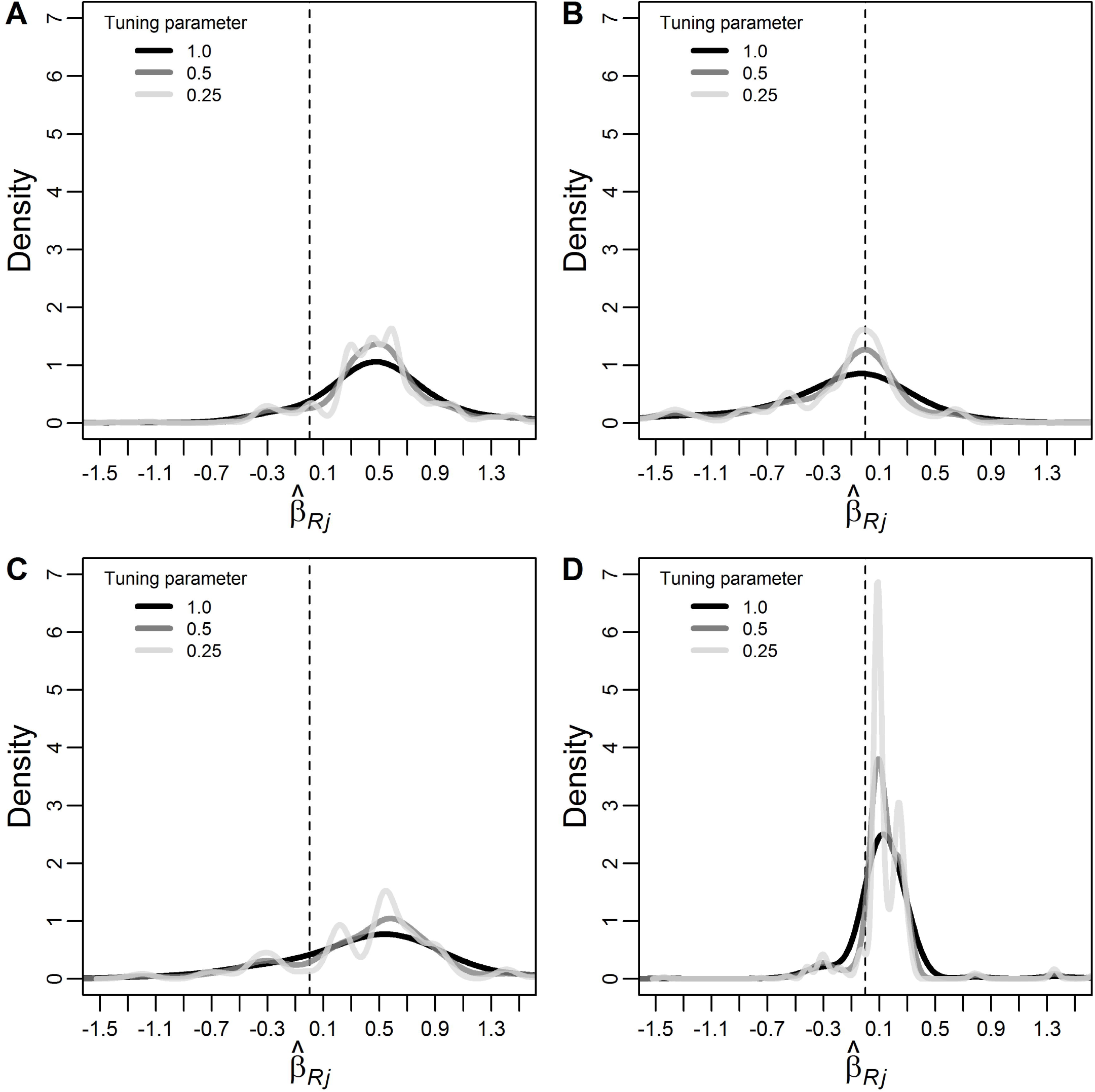
Weighted^a^ empirical density function of all individual-instrument ratio causal effect estimates 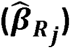 of plasma LDL-C (panel A), HDL-C (panel B), triglycerides (panel C) and urate (panel D) levels on coronary heart disease risk for different values of the tuning parameter ***φ***. LDL-C: low-density lipoprotein cholesterol. HDL-C: high-density lipoprotein cholesterol. The dashed line indicates the zero value. ^a^Weights were calculated without making the NOME assumption.

Results of the MR analysis are shown in Table 4. The smallest values of 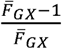 and 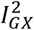 were 0.996 and 0.993, respectively, suggesting that IVW and MR-Egger regression estimates were not materially affected by regression dilution bias. P-values of the Cochran’s Q test ranged from 0.0003 (urate) to 1.7×10^−21^ (HDL-C), thus providing strong statistical evidence for heterogeneity between the ratio estimates. Nevertheless, results for LDL-C and triglycerides consistently suggested risk-increasing causal effects. In the case of HDL-C, the IVW method suggested a protective effect, with 1 standard deviation increase in HDL-C being associated with a 0.254 (95% CI: 0.115; 0.393) decrease in CHD ln(odds). However, the other methods did not confirm this result, suggesting that it was due to negative horizontal pleiotropy (as suggested by visually inspecting the distribution of ratio estimates). Finally, the IVW method suggested a 0.163 (95% CI: 0.027; 0.298) increase in CHD ln(odds) per standard deviation increase in urate levels. Other methods did not confirm this finding, suggesting that it could be a result of positive horizontal pleiotropy (as the empirical density plot suggested).

**Table 4.** Mendelian randomization estimates of the causal effect of urate plasma levels (in standard deviation units) on CHD risk (in ln(odds)) using 31 genetic instruments.

| Exposure | Estimator | Beta | SE | 95% CI | P-value |
| --- | --- | --- | --- | --- | --- |
| LDL-C | IVW | 0.476 | 0.060 | 0.357; 0.595 | $1.8 \times 10^{-11}$ |
| | MR-Egger - $\beta_0$ | -0.009 | 0.005 | -0.020; 0.001 | 0.083 |
| | MR-Egger - $\beta_1$ | 0.624 | 0.103 | 0.419; 0.828 | $5.3 \times 10^{-8}$ |
| | Weighted median | 0.457 | 0.064 | 0.331; 0.583 | $7.4 \times 10^{-10}$ |
|  | Simple MBE<sup>a</sup> | 0.422 | 0.187 | 0.056; 0.788 | 0.027 |
| | Weighted MBE<sup>a,b</sup> | 0.491 | 0.109 | 0.276; 0.705 | $2.7 \times 10^{-5}$ |
| HDL-C | IVW | -0.254 | 0.070 | -0.393; -0.115 | $4.9 \times 10^{-4}$ |
| | MR-Egger - $\beta_0$ | -0.014 | 0.005 | -0.025; -0.003 | 0.011 |
| | MR-Egger - $\beta_1$ | -0.013 | 0.115 | -0.241; 0.215 | 0.913 |
|  | Weighted median | -0.069 | 0.068 | -0.202; 0.065 | 0.314 |
|  | Simple MBE<sup>a</sup> | -0.174 | 0.171 | -0.509; 0.161 | 0.311 |
|  | Weighted MBE<sup>a,b</sup> | -0.003 | 0.088 | -0.175; 0.170 | 0.974 |
| Triglycerides | IVW | 0.416 | 0.081 | 0.252; 0.580 | $6.0 \times 10^{-6}$ |
| | MR-Egger - $\beta_0$ | 0.000 | 0.007 | -0.015; 0.015 | 0.962 |
| | MR-Egger - $\beta_1$ | 0.422 | 0.140 | 0.140; 0.704 | 0.004 |
| | Weighted median | 0.516 | 0.083 | 0.352; 0.679 | $1.5 \times 10^{-7}$ |
|  | Simple MBE<sup>c</sup> | 0.875 | 0.259 | 0.367; 1.383 | 0.002 |
| | Weighted MBE<sup>c,b</sup> | 0.547 | 0.134 | 0.284; 0.810 | $1.8 \times 10^{-4}$ |
| Urate levels | IVW | 0.163 | 0.066 | 0.027; 0.298 | 0.020 |
| | MR-Egger - $\beta_0$ | 0.008 | 0.005 | -0.002; 0.018 | 0.118 |
| | MR-Egger - $\beta_1$ | 0.048 | 0.096 | -0.148; 0.245 | 0.614 |
|  | Weighted median | 0.119 | 0.061 | -0.001; 0.239 | 0.061 |
|  | Simple MBE<sup>c</sup> | 0.188 | 0.163 | -0.132; 0.507 | 0.259 |
|  | Weighted MBE<sup>c,b</sup> | 0.092 | 0.066 | -0.038; 0.221 | 0.175 |

LDL-C: low-density lipoprotein cholesterol. HDL-C: high-density lipoprotein cholesterol. IVW: Inverse-variance weighting. SE: Standard error. CI: confidence interval. MBE: mode-based estimate.
<sup>a</sup> $\varphi=0.5$ .
<sup>b</sup>Not under the NO Measurement Error (NOME) assumption.
<sup>c</sup> $\varphi=0.25$ .

## Discussion

We have proposed a new MR method – the MBE – for causal effect estimation using summary data of multiple genetic instruments. Its performance was evaluated in a simulation study and its application illustrated in a real data example. An overview of the summary data MR methods that we evaluated (as well as the simple median) is provided in Table 5.

**Table 5.** Breakdown level and assumptions regarding horizontal pleiotropy of the inverse variance weighted (IVW), MR-Egger regression, simple and weighted median, and simple and weighted MBE’s.

| Method | Breakdown level | Assumptions regarding horizontal pleiotropy |
| --- | --- | --- |
| IVW | 0% | Consistent if the sum of horizontal pleiotropic effects of all instruments is zero. |
| MR-Egger regression | 100% | Consistent even if all instruments are invalid if InSIDE holds. |
| Simple median | $100\left(\frac{\lfloor L/2 \rfloor + 1}{L}\right)\%$ | Consistent if less than 50% of instruments are invalid, regardless of the type of horizontal pleiotropy. |
| Weighted median | 50% (exclusive) | Consistent if less than 50% of the weight is contributed by invalid instruments, regardless of the type of horizontal pleiotropy. |
| Simple MBE | Ranges from $100\left(\frac{\lfloor L/2 \rfloor + 1}{L}\right)\%$ to $100\left(\frac{L-2}{L}\right)\%$ | Consistent if the most common horizontal pleiotropy value is zero (i.e., ZEMPA), regardless of the type of horizontal pleiotropy. |
| Weighted MBE | Ranges from 50% (exclusive) to 100% (exclusive) | Consistent if the largest weights among the $k$ subsets are contributed by valid instruments (i.e., ZEMPA), regardless of the type of horizontal pleiotropy. |

IVW: Inverse-variance weighting. InSIDE: Instrument Strength Independent of Direct Effect. ZEMPA: ZERo Mode Pleiotropy Assumption. MBE: mode-based estimate.

Consistent causal effect estimation using the MBE requires that ZEMPA holds. ZEMA is an assumption that relates to the underlying bias parameters (the *b*_*j*_’s) that contribute to the ratio estimand *β*_*j*_ = *β* + *b*_*j*_ identified by the *j*th genetic instrument. If ZEMPA is satisfied then the MBE yields a consistent estimate for the causal effect. However, due to imprecision in the 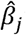 ‘s in finite samples, in practice the MBE may be contaminated by some invalid invariants even if ZEMPA holds. This can be seen in our simulations, where ZEMPA is only violated when all instruments are invalid, but nevertheless there is bias in the MBE when some of the instruments are valid. In practice, the MBE also depends on the magnitude of the bias, with invalid genetic instruments identifying causal effect parameters that are close to the true causal effect being more likely to contaminate the MBE estimate. However, this also means that genetic instruments that would introduce strong bias are less likely to contaminate the MBE.

In our simulations, we evaluated eight different versions of the MBE. Decreasing the tuning parameter *φ* reduced bias (at the cost of reduced precision) when horizontal pleiotropy does not violate the InSIDE assumption. However, when InSIDE is violated, a similar behaviour could only be clearly seen for the simple MBE. Choosing the value of the tuning parameter *φ* is a bias-variance trade-off and depends on how stringent the smoothing bandwidth needs to be and how stringent it can be before being prohibitively imprecise. In our applied example, we identified the stringency required through a graphical examination, and verified that the MBE’s were powered enough to detect a causal effect between HDL-C and triglycerides on CHD risk. Moreover, in the case of urate levels, the weighted MBE was similarly precise to that of the IVW and weighted median methods. This suggests that it may be feasible to set *φ* to very stringent values in practice, especially when there are multiple instruments selected based on genome-wide significance. Evaluating a range of *φ* values through a graphical examination may be useful to investigate how susceptible the MBE is to contamination from invalid instruments.

Assuming NOME increased bias and reduced the coverage of the 95% confidence intervals in the presence of invalid instruments, but reduced regression dilution bias and improved power in the two-sample setting. However, such gains were relatively small and virtually disappeared in simulations with larger sample sizes. Moreover, the results in the applied example were virtually identical whether or not NOME was assumed. These findings suggest that the NOME assumption is not necessary (and might be even unwarranted) when deriving weights for the MBE.

Although the simple MBE was less precise than the weighted MBE, it was less prone to bias due to violations of the InSIDE assumption. However, it was more prone to bias when InSIDE holds. Indeed, a similar pattern has been previously shown for the simple and weighted median. ^10^ This suggests that comparing both methods would be a useful sensitivity analysis in practice, although care must be taken since the simple MBE may in some cases (as in our real data example with urate levels) be prohibitively imprecise. Importantly, all the recommendations above are general, and we strongly encourage researchers to consider study-specific factors when deciding upon these aspects. One way of doing so is to perform simulations that reflect the study-specific context and compare different thresholds and filters in a range of different scenarios, keeping observable parameters (e.g., sample size) constant. Such simulations would also be useful to identify how strong the violations of the assumptions must be in order to obtain the observed results, which may be a useful sensitivity analysis that will either strengthen or weaken causal inference.

In our simulations, the 95% confidence intervals of the MBE computed using the normal approximation presented over-coverage (i.e., coverage larger than 95%). This may be due to the MBE being less influenced by outlying causal effect estimates (which is indeed the basis of the method), which correspond to the most imprecise ones when all instruments are valid. Therefore, the causal effect estimate fluctuates less around the true causal effect *β* (i.e., is less influenced by sampling variation). This may also explain the less pronounced over-coverage in the weighted median. We compared the normal approximation with the percentile method (Supplementary Table 7), but over-coverage in the latter was even greater when there were no or few invalid instruments. Moreover, after a certain proportion of invalid instruments (around 50%) coverage of the percentile method reduced markedly, while this occurred gradually in the normal approximation method. We therefore proposed the latter method to compute confidence intervals, but there might be better alternatives.

Another aspect of the MBE method (and of the weighted median) that requires further research is regression dilution bias in the two-sample setting. Dilution in the IVW estimate and MR-Egger regression estimates can be quantified by the 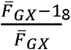 and 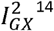 ^14^ statistics, respectively. Understanding how regression dilution bias operates in IVW and MR-Egger contributed to develop correction methods, thus reinforcing the importance of research in this area regarding the MBE and the weighted median methods.

Although this is the first description of using the MBE as a causal effect estimate in MR, other closely related methods have already been published. For example, Guo et al. ^22^ have recently described a method based on bivariate comparisons of all pairs of instruments, which classify instruments as estimating or not estimating the same causal effect. The largest identified set of concordant instruments can then be used to estimate the causal effect using, for example, the IVW method. Therefore, Guo et al.’s approach also relies on the assumption that the most common causal effect estimate is a consistent estimate of the true causal effect (i.e., ZEMPA). In fact, both our approach and Guo et al.’s can be viewed as methods that fully exploit the power of the consistency criterion defined originally by Kang et al, ^23^ who used it to propose a LASSO-based variable selection procedure to detect and adjust for pleiotropic variants. However, Guo et al.’s method and the MBE (which was developed independently from their work) are very different in their implementation. Ours is designed to be simple to understand and implement, does not require selecting instruments, and is easy to extend to any weighting scheme one desires. Moreover, plotting the empirical density function using different bandwidths may be a useful tool to visually explore the distribution of the 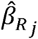s, and provides an intuitive way to select the optimal bandwidth value. In separate work we conduct a thorough review Guo et al.’s method after translating it to the two-sample context, and suggest some simple modifications to improve its performance. ^24^

It is also important to consider that there are other strategies to compute the mode of continuous data. In preliminary simulations, the modified Silverman’s rule was both generally more robust against horizontal pleiotropy than the original Silverman’s rule^25^ and more powered to detect a causal effect. Therefore, we opted for the modified rule. However, many other kernels and bandwidth selection rules could be used, as well as strategies that are not based on the smoothed empirical density function, such as the simple and robust parametric estimators, ^17^ Grenander’s estimators^26^ and the half-sample mode method.^16^ Further research is required to translate these mode estimators into the summary data MR context and compare their performance under different scenarios.

We propose the MBE as an additional MR method that should be used in combination with other approaches in a sensitivity analysis framework. Using several methods that make different assumptions, rather than a single method, is a useful strategy to assess the robustness of the results against violations of the instrumental variable assumptions^27,28^. Further developments in this area (including some aspects of the MBE itself) will contribute to expanding the arsenal of tools available to applied researchers to interrogate causal hypotheses with observational data.

## Supplementary Figure Legends

**Supplementary Figure 1. Histograms and smoothed empirical density plots of a hypothetical bimodal variable *V* (true mode is zero).**

